# Complex social behaviour during an extended period of time in a valproic acid animal model of autism spectrum disorder

**DOI:** 10.1101/2022.10.24.513470

**Authors:** Alexandre Maisterrena, Fabrice de Chaumont, Jean-Emmanuel Longueville, Eric Balado, Elodie Ey, Mohamed Jaber

**Affiliations:** Université de Poitiers, Inserm, Laboratoire de Neurosciences Expérimentales et Cliniques U1084, Poitiers, France; Institut Pasteur, CNRS, Gènes synapses et cognition, Paris, France; CNRS, Institut de Génétique et de Biologie Moléculaire et Cellulaire, Illkirch, France; CHU de Poitiers, Poitiers, France

**Author notes:** **Corresponding author.** Université de Poitiers, Inserm, Laboratoire de Neurosciences Expérimentales et Cliniques, Bâtiment B36, 1 rue Georges Bonnet, BP 633, TSA 51106, 86073 POITIERS cedex9 – FRANCE.

**Keywords:** autism spectrum disorder, machine learning, Live Mouse Tracker, social behavior, female, valproic acid, mouse models

## Abstract

Autism Spectrum Disorder (ASD) is a progressive neurodevelopmental disorder characterized mainly by deficits in social communication and stereotyped and restricted interests. Deficits in social interactions in ASD animal models are generally analysed using the three chamber’s test paradigm that is simple to implement and use but fails to detect subtle social deficits or complex social behavior on an extended period of time within a group of mice. Here, we set up a novel procedure entitled the Live Mouse Tracker (LMT) that detects a great number of complex social behaviours that we recorded continuously for up to three days in groups of 4 mice. This was performed in the valproic acid (VPA) mouse model where VPA (450 mg/kg) was injected to pregnant females at E12.5. Studies were performed with a special focus on females given that ASD is 3-4 times more diagnosed in males than in females and that several ASD models failed to detect major social deficits in females, contrary to males. Comparisons were made within groups of 4 female animals with same treatment or within groups of different treatments (saline versus VPA). We report that VPA females show several types of social deficits and that are different in nature and magnitude in relation with time (from 1 hour to 3 days). These deficits were also different when VPA mice were tested together compared to when they were mixed with saline treated mice. Indeed, while social behavior was improved in VPA mice with the presence of saline mice that of saline mice was negatively affected by the presence of VPA mice.

This study indicates that female VPA mice show several social deficits, contrary to the common knowledge. It further implies that ASD related behavior alters normal behavior in a mixed group of mice.

## Introduction

Autism spectrum disorder (ASD) is a pervasive developmental disorder that is currently diagnosed based on two main clusters of behaviors as per the current version of the Manual of the American Psychiatric Association: (i) deficits in social communication and interactions and (ii) repetitive and restrictive behavior and thoughts. Symptoms need to be present since childhood and affect negatively the person’s life^1^. Symptoms and comorbidities vary in their expression and severity leading to a spectrum of disorders. ASD is one the leading developmental disorders with a significant burden on the affected person, its family and the society in general. Its prevalence is in constant increase probably due to a combination of factors^2^. These include a heightened awareness of the disease by family and practitioners leading to a better diagnosis and recognition but also to possibly increased environmental insults from our modern way of living^3,4^. ASD has a prevalence of 3:1 in males to females although this proportion is still a matter of debate^5^. Several hypotheses are associated with this sexual dimorphism, among which is the possibility that females are underdiagnosed as ASD symptoms may be expressed differently in relation to gender^5^.

The specific causes of ASD are still not clearly identified but there is a general consensus that there is no single etiology, rather a combination of both genetic and environmental factors^6^. There are over 1000 genes that have been implicated in ASD, each being responsible for only a minor proportion^7^. A common ground for most of these genes is that they seem to be implicated in neuronal transmission with a great part being at the interface of glutamatergic and GABAergic synapses^8–10^. Gene mutations all together are thought to account for only 25% of ASD cases so that environmental factors may account for the rest. Effects of these environmental factors are most inflicting if occurring during prenatal development and include toxins, pharmacological agents and viral and bacterial infections^11^.

Knowledge of genetic and environmental determinants of ASD has led to the development of various genetic and environmental mouse models mimicking the main ASD symptomatology and thus showing good construct and face validity for most of them, or at least striving towards that direction^12,13^. Among these animal models, the valproic acid (VPA) administration as a single dose (250-500 mg/kg) to pregnant females at a critical age of embryonic development (E 12.5) has been constantly shown to be a reliable ASD model recapitulating several behavioral, cellular and molecular phenotypes of the disorder^13–18^.

As stated above, social deficits are amongst the hallmark symptoms of ASD and have been reported in these ASD models using various procedures, the most common one being the 3-Chamber test (3-CT) apparatus that was first set in place by Crawley in 1999^19^. This is a simple test measuring a mouse social behavior in a small surface subdivided into three chambers and counting the time, in frames of 10 minutes, that a mouse spends in a chamber containing another mouse compared to an empty chamber. If a new animal is thereafter introduced, one can also measure the preference to social novelty of the analyzed mouse, a behavior that is also affected in ASD patients. The 3-CT and similar procedures, including measuring social behavior in an open field, are widely used amongst researchers dealing with social determinants of ASD; they have the benefits of being easy to set in place and yield straightforward scores to analyze statistically^20^. As such, they are very useful for initial screenings but do not permit the analyses of complex social behavior, during an extended period of time, within a group of mice.

Here, we have implemented the recently described Life Mouse Tracker (LMT) procedure that enables automatic live tracking and subsequent analyses of complex social behavior in a group of mice with virtually no time limit but the one set by the experimenter^21^. This is achieved through the combination of computer vision through a depth-sensing infrared camera combined with machine learning and radio frequency identifying chips for animal identification. The social profile of each animal can thus be obtained, and its individual and group behavior analyzed in depth. Currently, this procedure must be set up starting from its basic hardware components by the experimenter. The corresponding software is in open access and can be downloaded directly but must be adapted to each’s needs.

Using this procedure, we analyzed a wide range of collective and individual social behaviors continuously for 3 days in groups of female mice that were prenatally exposed to either a saline solution or VPA (450 mg/kg) at E12.5. Analysis was performed on groups of 4 mice that were either homogeneous or mixed regarding their treatment. Our study provides a wide scale behavioral analysis of several parameters related to social behavior at different periods of time ranging from 1hr to 3 days and reveals a wide spectrum of deficits in ASD mouse models that can also affect control mice when housed together.

## Materials and Methods

### Hardware- and software-related information

The LMT equipment needs to be built from scratch by the user according to the previously described procedure as no commercial version is yet available^21^. This system enables individual identification of groups of mice over several days, their automatic live tracking using a random-forest machine learning process for animal and posture identification, radio-frequency identification (RFID) chips detected by floor antennas for mice identification, and an infrared-depth RGB-D camera. For this, a Kinect camera is set 63 cm above a 50 × 50 cm2 cage, perpendicular to the floor. RFID antennas (134-125 kHz) are placed under the cage and induce an electric current in the RFID chips carried by a mouse thus allowing their identification. The depth map is represented as an image of 512 × 424 pixels and the segmentation map represents all the animals and objects in the field with a depth sensitivity equal to 1,4 mm.

Behavioural labelling is performed in real time using the continuous stream of depth and volume information to characterize the shape and posture of each individual, as well as the relative position between individuals. The segmentation process detects all moving objects (a mouse size corresponds to 60 pixels approx.) and filters small ones (less than 30 pixels). To obtain all individual segmentations, a connected component extraction is performed. An additional filter is set to determine that the moving objects are indeed mice, and this is achieved through a machine learning algorithm that is trained during the first seconds of the observations. A detection splitter recovers animals fused into a single segmentation, based on an index map that corresponds to the identity index of each pixel of the mask. Thus, the system determines the outline mask and orientation of each mouse and builds a comprehensive repertoire of individual and social behavioural events from these data.

At the end of the detection procedure, different processes are used to associate a detection with a track or identity. The feature vector for machine learning is computed for each detection and allows machine learning algorithm to classify objects; it is composed of 33 values, the first being the ID of the animal, 16 values are the infrared histogram, and the 16 others are the depth histogram. The feature vector will therefore act as the object signature. The machine learning algorithm is involved in preparing the data, searching for the identity in concurrence with RFIDs and post-processing detections for the orientation of the animal. Information on the detection, tracking and RFID readings of mice is stored in a database and can be freely accessed live or later as needed. Video and background maps are simultaneously recorded as videos and image series, respectively.

All software can be downloaded directly from the following site: http://www.bioimageanalysis.org/lmt/. Full source code is available at http://icy.bioimageanalysis.org/plugins/livemousetracker. This includes Java code and CAD hardware resource files. Python analysis scripts are available at https://github.com/fdechaumont/lmt-analysis.

### Animals

Animal housing and experimental procedures were performed in accordance with the European Union directive (2010/63/EU) and validated by the regional ethical committee (Approval # 2020022610505052, 2021011214447548 and 2022021015414724). C57BL/6Jrj Mice (Janviers labs, France) were housed at the local Prebios animal facility, in ventilated cages with access to food and water *ad libitum*. Room temperature was maintained at 23°C on a 12h light/dark cycle.

### Experimental Design

27 females and 22 males were used for mating. For this, 2 females were placed with a single male and left overnight, and males were retrieved from the cage following mating. Pregnant mice received a single i.p. injection of either VPA (450 mg/kg) or NaCl 0.9 % at gestational day 12,5 (E12.5) when the neuronal tube is closing in rodents, followed by neurogenesis and neuronal migration^17,22^. Following injection, pregnant mice were left undisturbed until they gave birth. At weaning on postnatal day 28 (P28), sex and age matched pups were separated and raised by groups of 4 in a randomized fashion to avoid littermate effects. Thus, LMT measurements were performed on groups of 4 mice that were not previously housed together. All mice were identified both by ear tags and with a RFID tag inserted subcutaneously under isoflurane anaesthetic with local analgesia (lidocaine 2% in ointment form). Female mice were then allocated to two experimental groups depending on prenatal treatment: VPA females (n=37), and saline females (n=37). The experiment timeline is presented in Figure 1. Behavioral analysis was performed for 3 continuous days for LMT and during 30 min within the light cycle for the 3-CT procedures. At the end of the behavioral experiments, mice were sacrificed, and brains harvested for further analysis as described below.

**Figure 1:**
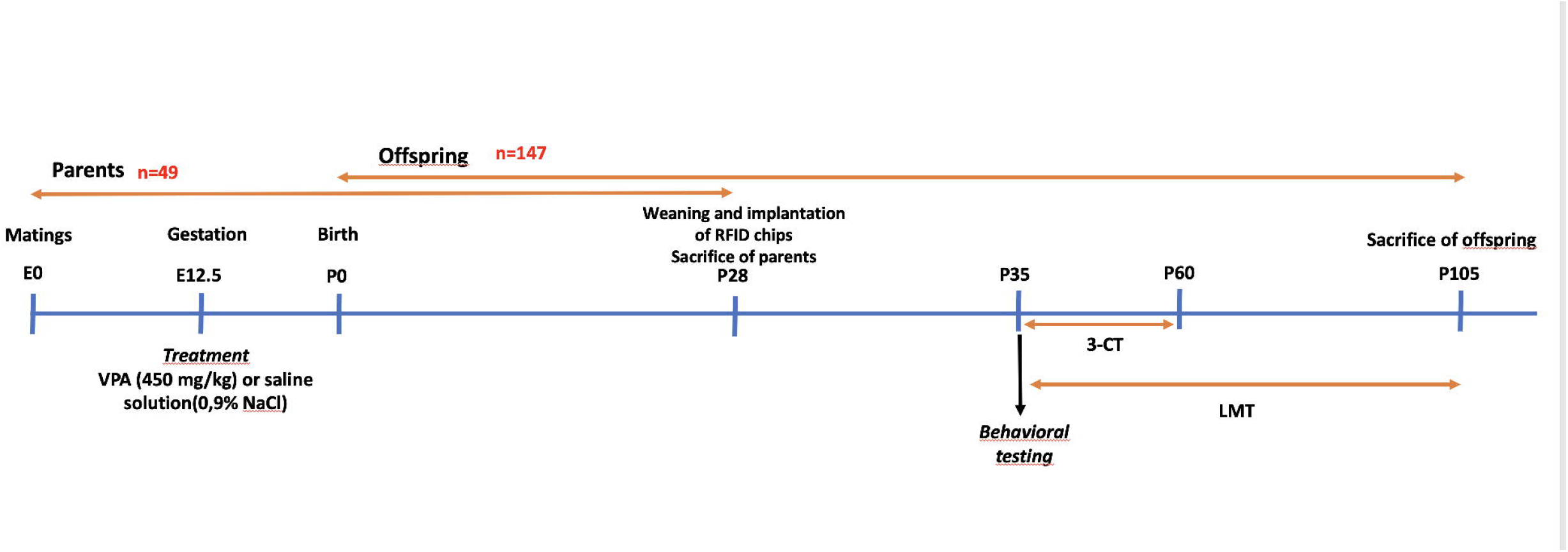
Experimental design and timeline. Pregnant female mice were i.p. injected at E12.5 with either saline (0.9% NaC) or a single dose of VPA (450 mg/kg). Offspring were weaned at 4 weeks pf age, implanted with a RFID tag and were allocated to 2 experimental groups in relation with prenatal exposure. Behavioral analysis was performed for 3 continuous days for the Live Mouse Tracker (LMT) and for 30 min. within the light cycle for the 3-Chamber test (3-CT) procedure.

### Individual social behavior as measured with the three chambers test (P30-P60)

Social interaction was assessed using the 3-CT as previously described^16^ on the VPA females (n=20), and saline females (n=15). The apparatus consists of a Plexiglas box (60×45×22 cm) partitioned into three chambers with retractable doorways. The first phase (PHASE-I) comprises two identical non-social stimuli (inverted wire-cups) placed in the opposite chambers. The second phase (PHASE-II) comprises a non-social stimulus and a social stimulus (a naïve age and sex-matched mouse with no previous contact with the tested mouse). Each phase was of 10 minutes during which time spent in each chamber and around the inverted cup was recorded using a video tracking system. Subsequently, a sociability index (SI) was calculated as follows: (time exploring social chamber – time exploring non-social chamber) / (time exploring social chamber + time exploring non-social chamber).

### Group behavior as measured with the Live Mouse Tracker (LMT) (P35-P105)

Data obtained through the LMT system served as the basis for computing several events that were inferred from shape geometry. Overall, 35 behavioural events were defined related to intrinsic and relative positions of the mice that were extracted automatically as previously described^21^. Behavioral features were split into five categories: (i) individual behavior, (ii) social dyadic events between two mice, (iii) dynamic events between two mice (approach, escape, following each other…), (iv) subgroup events with 2, 3 or 4 mice, (v) group making and group-braking events. Details of these events can be found in the initial report of LMT and are reported in the results section^21^. It is to be noted however, that all behavior from a given category does not yield values of equivalent scale. Thus, and for the sake of presentation, behaviors with similar scales were gathered within a single figure, independent of their behavior category.

### Data analysis

Data are expressed as mean ± Standard Error of the Mean (SEM) and analyzed using GraphPad Prism-7 software (La Jolla, California, USA). Data that followed a normal distribution was analyzed using one-way or two-way analysis of variance (ANOVAs) whenever appropriate. Upon significant main effects, Tukey’s or Fisher’s LSD multiple comparisons were performed for behavioral or histological measures, respectively. When data did not follow a normal distribution, we conducted non-parametric tests (Kruskal-Wallis or Mann-Whitney as indicated) followed by Dunn’s multiple comparison tests. For all analyses, a p value of <0,05 was considered significant. All detailed statistical analysis and raw data can be obtained upon request to the authors.

## Results

### Social interaction analysis using the three-chamber test

The 3-CT was used to assess individual social behavior of female mice prenatally exposed to VPA (450 mg/kg) or saline at E12.5 as previously described^16^. Using this procedure, we have previously shown that only male mice that were exposed prenatally to VPA showed a dramatic decrease (down 7-fold) in their social behavior where VPA females had normal sociability^16^.

Here, none of the treatment groups, regardless the sex, showed spontaneous preference for any of the chambers during the 10 minutes habituation (PHASE I) (Figure 2). Both saline and VPA females spent more time in the social chamber than in the empty chamber (62% *versus* 38%, and 69% versus 30%, respectively) [F(3, 64)=43, p<0,0001], although VPA mice seem to show an even better preference for the social chamber than the saline group (p<0.05) (Figure 2). However, when a female mouse was introduced in the opposite chamber, VPA female mice failed to show social novelty preference contrary to saline female mice (figure 2) [F(3, 64)=6,8, p<0.001]. Of interest is also the finding that saline mice preferred the new social chamber (SC2) more than the VPA mice (p<0.05).

**Figure 2:**
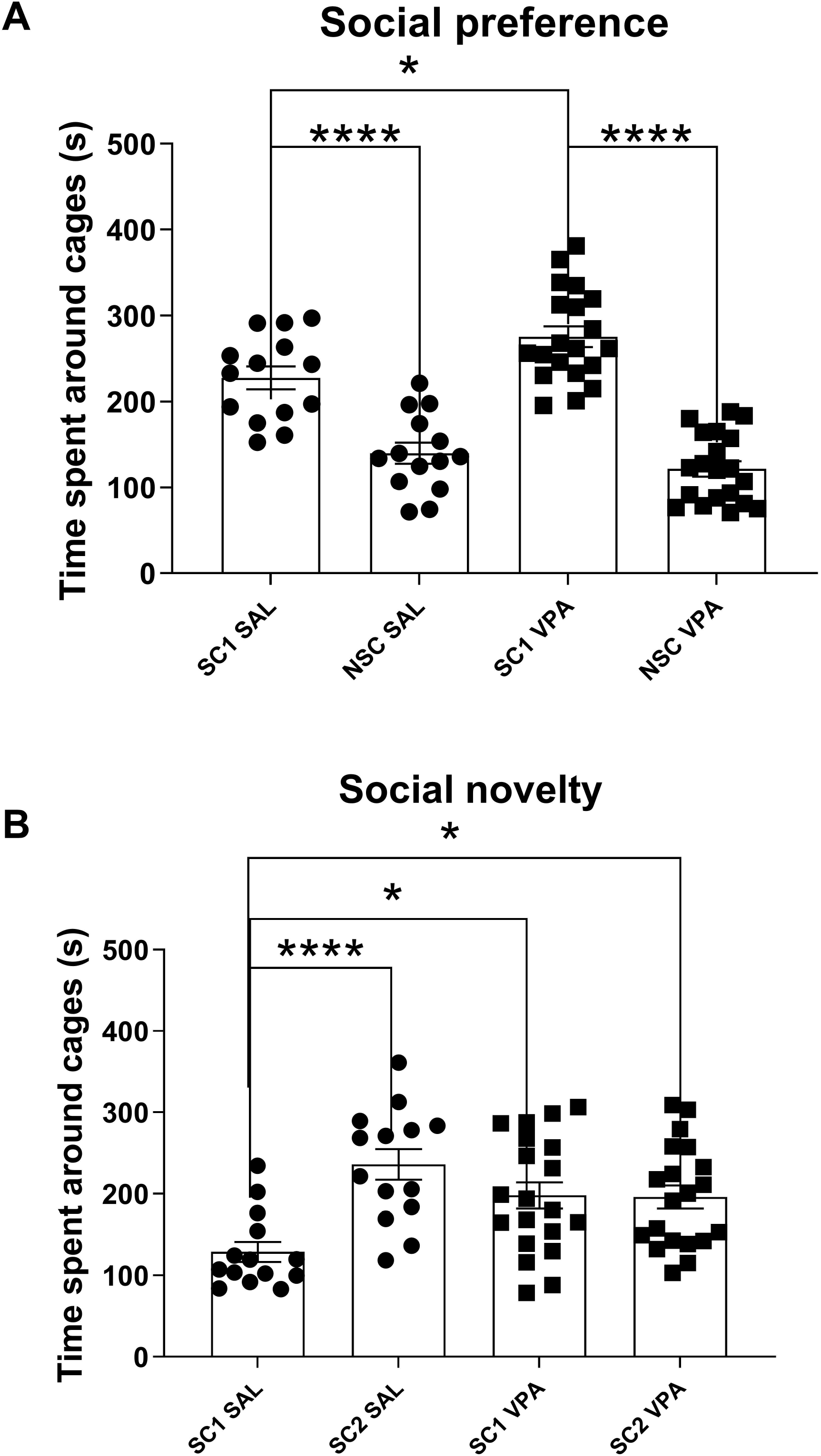
Social behavior as analyzed in the 3-chamber test (3-CT) procedure. A) Following habituation (Phase I) mice were put in 3-CT with one chamber containing an encaged mouse (social chamber 1: SC1) and one chamber with an empty cage (non-social chamber: NSC) to evaluate social preference (Phase II). (B) a novel mouse was introduced in the chamber 2 (social chamber 2: SC2) in addition to the encaged mouse in SC1 to evaluate social novelty recognition (Phase III). During Phase III, VPA female mice spent equal time interacting with SC1 and SC2 while saline mice spent more time interacting with SC2. All data are expressed as means ± SEM; One way ANOVA followed by Tukey’s multiple analysis was performed (*p<0.05, **p<0.01, ***p<0.001). Saline females (n=14), VPA females (n=20).

### Social interaction analysis using the Live Mouse Tracker system for 3 continuous days

We analyzed several variables of social behavior as described in the materials and methods section, for 3 days, on homogenous (same treatment within the group) or mixed groups (one VPA and three saline mice within the group) of 4 female mice that were prenatally (E 12.5) exposed to either saline or VPA (450 mg/kg). Mice behavior was segmented into 5 groups of behavior as described in the methods section and as initially proposed^21^. As reported in figure 3, few altered behaviors can be identified at the end of the three days period in VPA groups of mice compared to control, whether in males or females. Indeed, only a few social behaviors were affected by the VPA treatment. This is for instance the case for the number of following behavior and movement in contact with another mouse for which VPA females performed less than saline.

**Figure 3:**
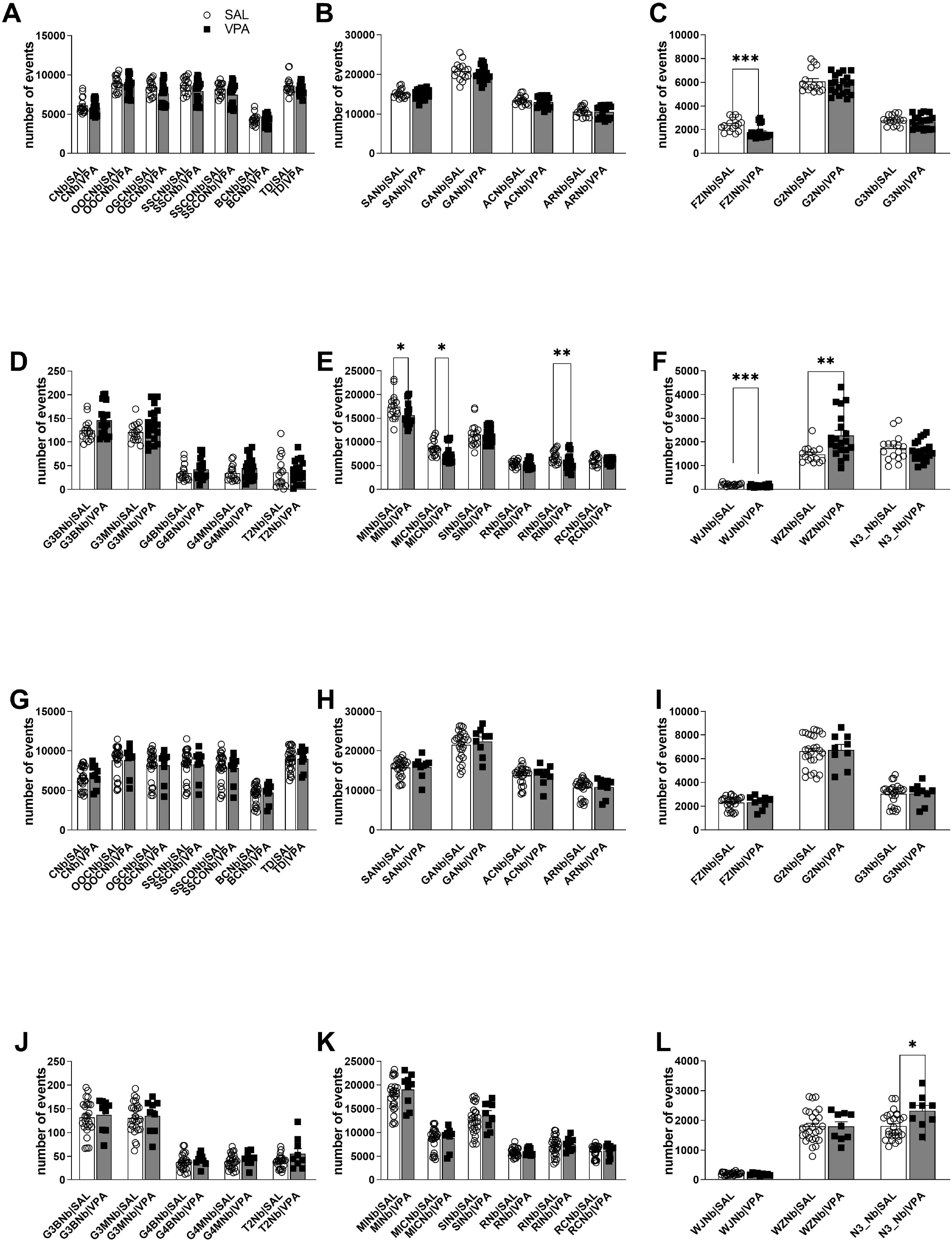
Social behavior during three continuous days as measured using the Life Mouse Tracker (LMT) procedure. (A) Number of various social contacts within the homogenous groups (mice of the same treatment) during the three days of experiment. (B) Number of various social approaches within the homogenous groups. (C) Number of “following behavior” and of clusters of 2 and 3 mice within the homogenous groups (D). Number of “making and breaking of groups” of clusters of 3 and 4 mice and number of “trains” of 2 mice within the homogenous groups. (E) Number of “moving behavior” and “rearing” within the homogenous groups. (F) Number of “wall jump”, “water zone interaction” and “nest of 3” within the homogenous groups. (G) Number of various social contacts for the mixed groups (3 saline and 1 VPA). (H) Number of various social approaches within the mixed groups. (I) Number of “following behavior” and of clusters of 2 and 3 mice within the mixed groups (J) Number of “making and breaking of groups” of clusters of 3 and 4 mice and number of “trains” of 2 mice within the mixed groups. (K) Number of “moving behavior” and “rearing” within the mixed groups. (L) Number of “wall jump”, “water zone interaction” and “nest of 3” within the mixed groups. All data are expressed as means ± SEM; Mann-Whitney test was performed (*p<0.05, **p<0.01, ***p<0.001). Saline females homogenous group (n=16); VPA females homogenous group (n=20); saline females mixed group (n=27); VPA female mixed group (n= 9).

We then performed the same analysis on heterogenous groups of mice of the same sex. For this, we introduced in the LMT arena one VPA mouse and three saline mice of the same sex and monitored their behavior for three continuous days. As shown in figure 3, we found that VPA mice created significantly more nests of 3 mice than the saline mice in the group (p<0,05). No other differences were found between the VPA and saline mice when analyzed together within a heterogeneous group.

Altogether, analysis of social behavior for a total of three days did not show major social deficits in the various groups of mice. This may be due to a flattening effect of behavioral deficiencies in the VPA group on a 3 days scale. In order to test this hypothesis, we underwent a further analysis of the data by time segments as detailed below.

### Social interaction analysis using the LMT on various time periods

Group social interactions can vary greatly in time. Analyzing results on the total time period of three days may mask major differences in the behavior of different groups and that may be due to reaction to novelty, whether environmental or social. In order to determine the behavior of homogenous and mixed groups of female mice, we further analyzed the data obtained from the LMT system and segmented it to periods of 1h, 3h and 1 day (Figure 4).

**Figure 4:**
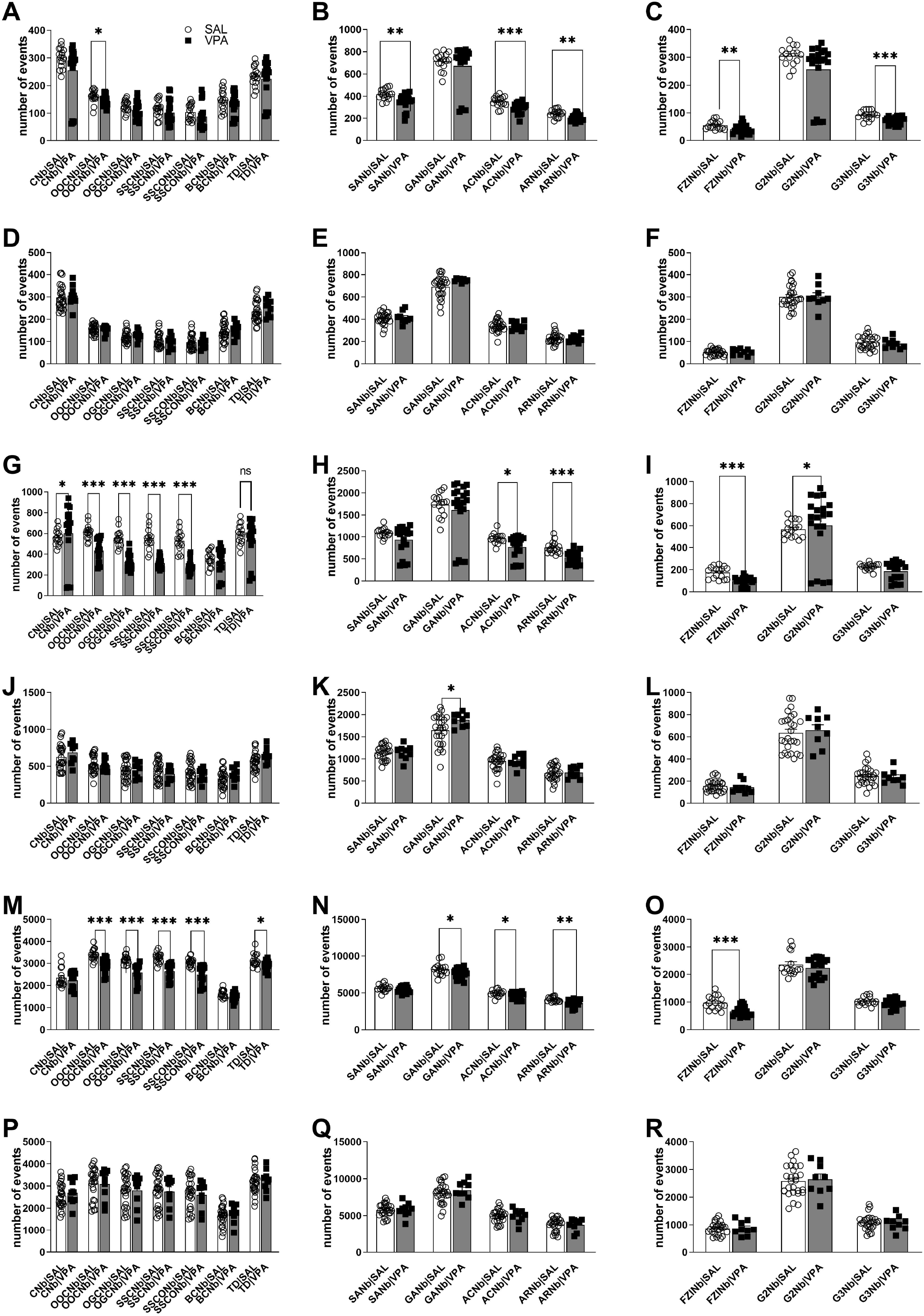
Social behavior at various time periods (1h, 3h and 12h) as measured using the Life Mouse Tracker (LMT) procedure. Number of various social contacts (A), social approaches (B) or of “following” and making groups of 2 and 3 (C) within the homogenous groups for 1 hour. Number of various social contacts (D), social approaches (E) or of “following” and making groups of 2 and 3 (F) within the mixed groups for 1 hour. Number of various social contacts (G), social approaches (H) or of “following” and making groups of 2 and 3 (I) within the homogenous groups for 3 hours. Number of various social contacts (J), social approaches (K) or of “following” and making groups of 2 and 3 (L) within the mixed groups for 3 hours. Number of various social contacts (M), social approaches (N) or of “following” and making groups of 2 and 3 (O) within the homogenous groups for 24 hours. Number of various social contacts (P), social approaches (Q) or of “following” and making groups of 2 and 3 (R) within the mixed groups for 24 hours. All data are expressed as means ± SEM; Mann-Whitney test was performed (*p<0.05, **p<0.01, ***p<0.001).

We report here that after one hour of experiment, there were few differences in the number of contacts between saline and VPA mice in both the homogeneous and the mixed groups (figure 4A and D). For example, VPA female mice of the homogenous groups performed significantly fewer social approaches than saline mice (−12% for SA versus saline, p<0.01) (figure 4B). No such differences were identified within the mixed groups (figure 4E). After 3h of experimentation, several deficits were identified in the number of social contacts within the VPA mice of the homogenous groups (−27% for oral-oral contacts p<0.0001; -42% for oral-genital contacts p<0.0001) (figure 4G) and some deficits in social approaches were also noticeable (−10% approach to contact, p<0.05) (figure 4H). Again, no major difference was found between saline and VPA mice in the mixed groups (figure 4J-L). We then analyzed the behavior after 24hours of experimentation. Again, we observed several deficits in the number of social contacts in the VPA mice in the homogenous groups (−11% oral-oral contacts p<0.0001; -13% oral-genital contacts, p<0.0001) (figure 4M), along with a slight reduction in the number of social approaches (−5% approach to contact, p<0.05) (figure 4N). Once more, no major differences were found in the mixed groups (figure 4P-R).

### Evolution in time of social behavior using the LMT

When analyzing social behavior in relation with time, we have identified major differences on individual and group behaviors. To further reveal these differences, we plotted social behavior of several chosen variables in relation with time (figure 5). We show here that across behaviors, the main differences between groups and treatment are observed during the first 24h to 48h of the experiment. For instance, we found significant alterations in the behavior of saline mice within the heterogenous groups during the first 24h (figure 5A). this was also the case in the VPA mice in the homogenous groups (figure 5A). Across time, differences between saline mice in homogenous groups (all saline) compared to the heterogenous groups (3 saline and one VPA mice) were observed but that were of a lesser magnitude than differences between saline and VPA mice in homogenous groups (figure5B-D).

**Figure 5:**
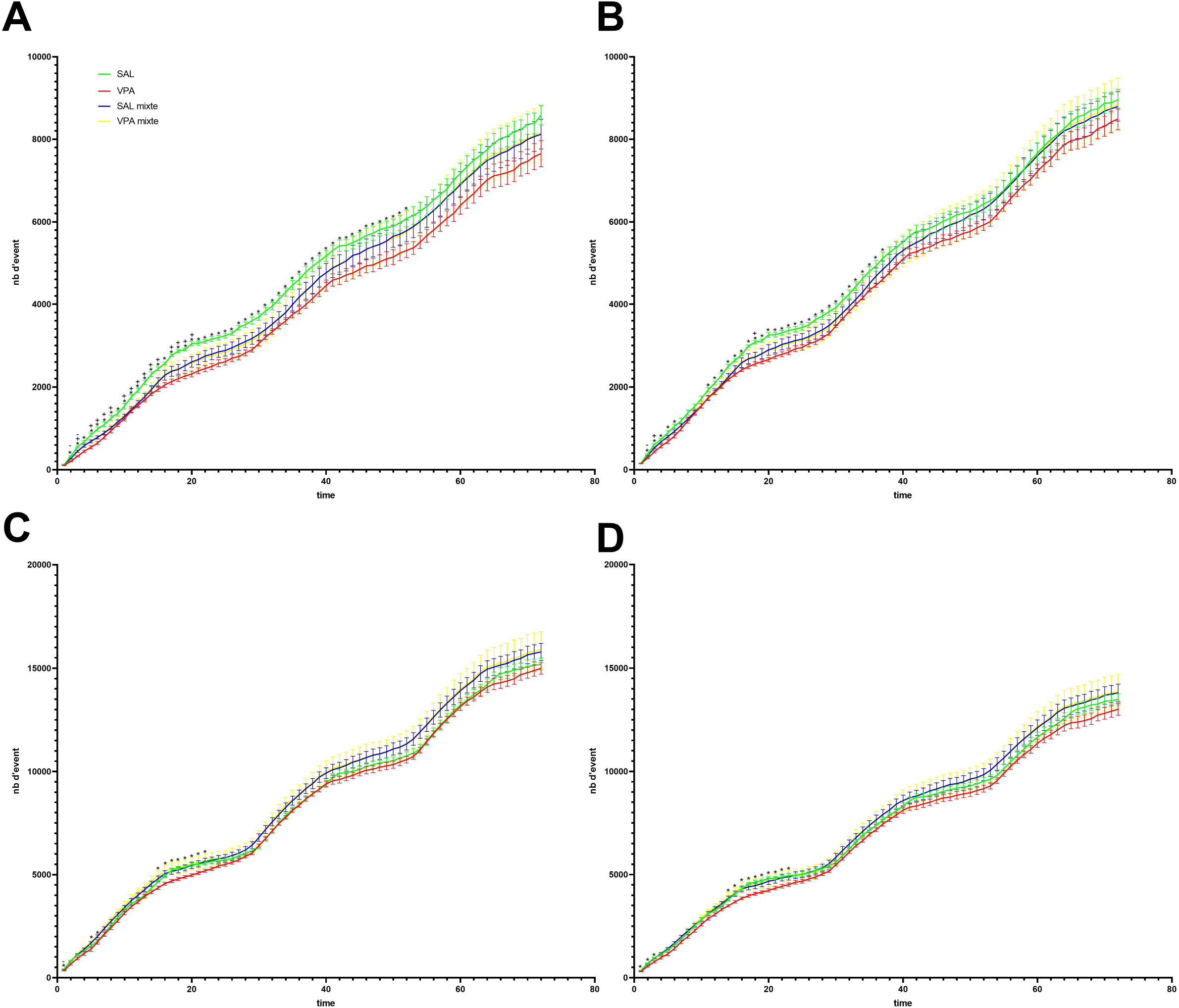
Evolution in time of social behavior using the Life Mouse Tracker (LMT) procedure. Number of (A) oral-genital contacts, (B) oral-oral contacts, (C) social approaches, (D) social contacts over the three days of the experiment. All data expressed as means ± SEM; two-Way ANOVA with post-hoc Tukey was performed. Only selected differences were highlighted for the sake of clarity (*: differences between saline homogenous and VPA homogenous mice; +:differences between saline homogenous mice and saline mixed groups mice; -: differences between VPA homogenous and VPA mixed groups mice).

## Discussion

Social deficits are one the two hallmarks of ASD symptoms as defined by the American Manual of Psychiatry in its latest version^1^. As face validity is key in animal modeling of diseases and disorders, it has become mandatory for ASD animal models, whether genetic or environmental, to express some forms of social deficits before being considered for further analysis and characterization^19,20,23–25^.

Social deficits in ASD children are expressed in the form of social play behavior deficits and lack of non-verbal social gestures such as pointing, showing, and giving^26–29^. Mice are very social mammals and exhibit a wide range of social behavior such as direct cross interactions, sniffing, following, regrouping, collective nesting, maternal care, general play behavior and sexual approaches^30–32^. A significant number of reports have described social impairments in a large majority of ASD animal models, including the VPA models^13,17,26^.

Since its initial implementation, the 3-CT has become rapidly the gold standard to measure social approach, social preference and, more generally, exploratory locomotion^31^. The procedure is now automated and is now widely used, in its original form or with slight variations, amongst teams working on various animal models of ASD^33^. This automated test was referred to by the team as being a “basic protocol for a standardized, high throughput social approach test for assaying mouse sociability“ and that “allows to quantify direct social approach behaviors when a subject mouse is presented with the choice of spending time with either a novel mouse or a novel object”^33^. Despite the many advantages of this simple and efficient procedure, the 3-CT suffers from several drawbacks as it cannot evaluate complex social behaviors, in a group of mice, and for a long period of time.

Recently, we have published a series of studies on 3 different ASD animal models, genetic (Shank3) and environmental (VPA and poly I:C) all showing major deficits in social behavior that were almost exclusively present in male ASD mice but not in females ^13,16,16,34^. These deficits were also variable in their magnitude and expression in relation with ASD’s etiology, and they include deficits in social preferences and/or social novelty. However, both males and females showed abnormal gait and motor coordination behaviors and correlative loss of cerebellar Purkinje cells. This raised the question of whether ASD females were somehow protected from contracting social deficits, whatever the disorder’s etiology, or whether the procedure that is used to evaluate social behavior is not sensitive enough to detect mild or specific social deficits. Of interest is the fact that loss of PC cells was restricted to different cerebellar subregions between males and females, and this may possibly underly differences in the behavioral expression of ASD in relation with mice gender^13^.

Here, we aimed to explore whether we can identify complex social deficits in female VPA mice and determine the extent and the nature of these social deficits. For this, we have used in the widely used VPA ASD animal model and that we have previously characterized extensively in our experimental settings, where VPA is injected prenatally to the pregnant female at E12.5 at 450 mg/kg in a single i.p. dose. We implemented the recently developed LMT apparatus and performed the experiments on female mice. We measured social behavior on groups of 4 mice randomized from different litter of the same sex and of the same treatment (homogeneous groups) or with a mixed treatment of saline and VPA. Experiments were performed for three consecutive days and data was analyzed following 1hr, 3 hrs, 24 hrs and three days.

VPA, is a potent antiepileptic drug^35^ and mood stabilizer^36^ that modulates GABA neurotransmission^37^ and regulates gene expression through inhibition of histone deacetylase (HDAC) activity^38^. It has been known since 1991 that VPA is teratogenic when administered during pregnancy^39^ and later it was consistently reported that it also significantly increases the risk of developmental delays including ASD^40^. Prenatal exposure to a single dose of VPA around the date of E12.5 is now widely used to induce in offspring the two cardinal ASD symptoms, social deficits and stereotyped behavior. The VPA model is considered a reliable model bearing a very strong construct and face validity^17,18,26^. In our hands, we found that the VPA animal model was the most robust of all three models that we had explored in our experimental settings (Shank 3 ΔC or prenatal exposure to either Poly I:C or VPA). Indeed, we previously reported that VPA mice showed significant deficits in social interactions along with major gait and motor disorders that were even detected at early perinatal ages^16^. Of interest is also the fact that social deficits were directly correlated in their severity with motor and gait deficits and also with loss of PCs within restricted cerebellar areas.

We report here that: (i) VPA females show several forms of social deficits as compared to saline females; (ii) social deficits were complex in their nature as they implicated several behavioral traits and variables such as approaches, following, direct interactions, breaking a group of mice and nesting ; iii) these social deficits appear early on during the experiments then tend to disappear within three days implicating possibly a reaction to novelty; (iv) one of the most intriguing findings that we report here is that VPA mice, when placed within a group of saline mice, show significant improvement in several social variables ; (v) conversantly, saline mice within mixed groups showed degradations of several variables linked to social behavior in the presence of a VPA mouse. In other words, placing an ASD mouse within a group of normal mice improves the social behavior of the VPA mouse but deteriorates that of normal mice. These findings need to be confirmed with further studies in ASD animal models and more importantly in humans, as they may have repercussions regarding the current policies of integrating ASD patients within groups of typical individual. A such, these results may also suggest that some form of follow up and companionship is needed to increase the chances of success of including ASD patients in regular schools. Although research reports on this subject are very scarce, it was shown that when intervention therapies are implemented in schools, ASD children had significantly higher social network inclusion and received more friendship nominations than ASD children without such an intervention^41,42^.

To our estimate, the 3-CT should always be the default procedure to use for screening social behavioral deficits in animal models of ASD or other psychiatric illnesses. The LMT is to be used when precise and detailed exploration of social deficits are needed as is the case with females who do not show such deficits in the 3-CT whatever the ASD etiology. The LMT is also useful to study long term social behavior in a homogeneous or heterogeneous groups of mice. Currently, the LMT does not allow measurements of other social behaviors such as mating or fighting. Surely, with the current developments, this may be overcome in the future.

## Supporting information

List of abbreviations in figures

## Acknowledgments

The research leading to the results detailed in this manuscript received recurrent funding from Poitiers university and Inserm. Authors are grateful for Emilie Dugast and Anais Balbous for help with the animal experimentation authorization documents, and the Prebios animal facility staff for help and dedication.

## Statements and Declarations

Authors declare no financial or non-financial interests that are directly or indirectly related to the work submitted for publication.

## References

1. American Psychiatric Association (2013). Diagnostic and Statistical Manual of Mental Disorders Fifth Edition. (American Psychiatric Association) 10.1176/appi.books.9780890425596.

2. Järbrink, K., and Knapp, M. (2001). The Economic Impact of Autism in Britain. Autism 5, 7–22. 10.1177/1362361301005001002.

3. Maenner, M.J., Shaw, K.A., Baio, J., EdS Washington, A., Patrick, M., DiRienzo, M., Christensen, D.L., Wiggins, L.D., Pettygrove, S., et al. (2020). Prevalence of Autism Spectrum Disorder Among Children Aged 8 Years - Autism and Developmental Disabilities Monitoring Network, 11 Sites, United States, 2016. MMWR Surveill Summ 69, 1–12. 10.15585/mmwr.ss6904a1.

4. Hertz-Picciotto, I., and Delwiche, L. (2009). The Rise in Autism and the Role of Age at Diagnosis. Epidemiology 20, 84–90. 10.1097/EDE.0b013e3181902d15.

5. Loomes, R., Hull, L., and Mandy, W.P.L. (2017). What Is the Male-to-Female Ratio in Autism Spectrum Disorder? A Systematic Review and Meta-Analysis. J Am Acad Child Adolesc Psychiatry 56, 466–474. 10.1016/j.jaac.2017.03.013.

6. Chaste, P., and Leboyer, M. (2012). Autism risk factors: genes, environment, and gene-environment interactions. Dialogues Clin Neurosci 14, 281–292.

7. Vorstman, J.A.S., Parr, J.R., Moreno-De-Luca, D., Anney, R.J.L., Nurnberger Jr, J.I., and Hallmayer, J.F. (2017). Autism genetics: opportunities and challenges for clinical translation. Nat Rev Genet 18, 362–376. 10.1038/nrg.2017.4.

8. De Rubeis, S., He, X., Goldberg, A.P., Poultney, C.S., Samocha, K., Cicek, A.E., Kou, Y., Liu, L., Fromer, M., Walker, S., et al. (2014). Synaptic, transcriptional and chromatin genes disrupted in autism. Nature 515, 209–215. 10.1038/nature13772.

9. Wang, W., Li, C., Chen, Q., van der Goes, M.-S., Hawrot, J., Yao, A.Y., Gao, X., Lu, C., Zang, Y., Zhang, Q., et al. (2017). Striatopallidal dysfunction underlies repetitive behavior in Shank3-deficient model of autism. J. Clin. Invest. 127, 1978–1990. 10.1172/JCI87997.

10. Guang, S., Pang, N., Deng, X., Yang, L., He, F., Wu, L., Chen, C., Yin, F., and Peng, J. (2018). Synaptopathology Involved in Autism Spectrum Disorder. Front. Cell. Neurosci. 12, 470. 10.3389/fncel.2018.00470.

11. Bölte, S., Girdler, S., and Marschik, P.B. (2019). The contribution of environmental exposure to the etiology of autism spectrum disorder. Cell Mol Life Sci 76, 1275–1297. 10.1007/s00018-018-2988-4.

12. Bey, A.L., and Jiang, Y. (2014). Overview of mouse models of autism spectrum disorders. Curr Protoc Pharmacol 66, 5.66.1-26. 10.1002/0471141755.ph0566s66.

13. Thabault, M., Turpin, V., Maisterrena, A., Jaber, M., Egloff, M., and Galvan, L. (2022). Cerebellar and Striatal Implications in Autism Spectrum Disorders: From Clinical Observations to Animal Models. IJMS 23, 2294. 10.3390/ijms23042294.

14. Rodier, P.M., Ingram, J.L., Tisdale, B., Nelson, S., and Romano, J. (1996). Embryological origin for autism: Developmental anomalies of the cranial nerve motor nuclei. J. Comp. Neurol. 370, 247–261. 10.1002/(SICI)1096-9861(19960624)370:2<247::AID-CNE8>3.0.CO;2-2.

15. Schneider, T., and Przewłocki, R. (2005). Behavioral Alterations in Rats Prenatally Exposed to Valproic Acid: Animal Model of Autism. Neuropsychopharmacol 30, 80–89. 10.1038/sj.npp.1300518.

16. Al Sagheer, T., Haida, O., Balbous, A., Francheteau, M., Matas, E., Fernagut, P.-O., and Jaber, M. (2018). Motor Impairments Correlate with Social Deficits and Restricted Neuronal Loss in an Environmental Model of Autism. Int J Neuropsychopharmacol 21, 871–882. 10.1093/ijnp/pyy043.

17. Nicolini, C., and Fahnestock, M. (2018). The valproic acid-induced rodent model of autism. Experimental Neurology 299, 217–227. 10.1016/j.expneurol.2017.04.017.

18. Chaliha, D., Albrecht, M., Vaccarezza, M., Takechi, R., Lam, V., Al-Salami, H., and Mamo, J. (2020). A Systematic Review of the Valproic-Acid-Induced Rodent Model of Autism. Dev Neurosci 42, 12– 48. 10.1159/000509109.

19. Crawley, J.N. (2004). Designing mouse behavioral tasks relevant to autistic-like behaviors. Ment Retard Dev Disabil Res Rev 10, 248–258. 10.1002/mrdd.20039.

20. Silverman, J.L., Yang, M., Lord, C., and Crawley, J.N. (2010). Behavioural phenotyping assays for mouse models of autism. Nat. Rev. Neurosci. 11, 490–502. 10.1038/nrn2851.

21. de Chaumont, F., Ey, E., Torquet, N., Lagache, T., Dallongeville, S., Imbert, A., Legou, T., Le Sourd, A.-M., Faure, P., Bourgeron, T., et al. (2019). Real-time analysis of the behaviour of groups of mice via a depth-sensing camera and machine learning. Nat Biomed Eng 3, 930–942. 10.1038/s41551-019-0396-1.

22. Morriss-Kay, G., Ruberte, E., and Fukiishi, Y. (1993). Mammalian neural crest and neural crest derivatives. Annals of Anatomy - Anatomischer Anzeiger 175, 501–507. 10.1016/S0940-9602(11)80209-8.

23. Haratizadeh, S., Parvan, M., Mohammadi, S., Shabani, M., and Nozari, M. (2021). An overview of modeling and behavioral assessment of autism in the rodent. Int. j. dev. neurosci. 81, 221–228. 10.1002/jdn.10096.

24. Takumi, T., Tamada, K., Hatanaka, F., Nakai, N., and Bolton, P.F. (2020). Behavioral neuroscience of autism. Neuroscience & Biobehavioral Reviews 110, 60–76. 10.1016/j.neubiorev.2019.04.012.

25. Ellegood, J., and Crawley, J.N. (2015). Behavioral and Neuroanatomical Phenotypes in Mouse Models of Autism. Neurotherapeutics 12, 521–533. 10.1007/s13311-015-0360-z.

26. Tartaglione, A.M., Schiavi, S., Calamandrei, G., and Trezza, V. (2019). Prenatal valproate in rodents as a tool to understand the neural underpinnings of social dysfunctions in autism spectrum disorder. Neuropharmacology 159, 107477. 10.1016/j.neuropharm.2018.12.024.

27. Hobson, J.A., Hobson, R.P., Malik, S., Bargiota, K., and Caló, S. (2013). The relation between social engagement and pretend play in autism: Social engagement and play in autism. British Journal of Developmental Psychology 31, 114–127. 10.1111/j.2044-835X.2012.02083.x.

28. Jordan, R. (2003). Social play and autistic spectrum disorders: a perspective on theory, implications and educational approaches. Autism 7, 347–360. 10.1177/1362361303007004002.

29. Rutherford, M.D., Young, G.S., Hepburn, S., and Rogers, S.J. (2007). A Longitudinal Study of Pretend Play in Autism. J Autism Dev Disord 37, 1024–1039. 10.1007/s10803-006-0240-9.

30. Panksepp, J., Siviy, S., and Normansell, L. (1984). The psychobiology of play: theoretical and methodological perspectives. Neurosci Biobehav Rev 8, 465–492. 10.1016/0149-7634(84)90005-8.

31. Ricceri, L., Moles, A., and Crawley, J. (2007). Behavioral phenotyping of mouse models of neurodevelopmental disorders: Relevant social behavior patterns across the life span. Behavioural Brain Research 176, 40–52. 10.1016/j.bbr.2006.08.024.

32. Vanderschuren, L.J.M.J., Achterberg, E.J.M., and Trezza, V. (2016). The neurobiology of social play and its rewarding value in rats. Neurosci Biobehav Rev 70, 86–105. 10.1016/j.neubiorev.2016.07.025.

33. Yang, M., Silverman, J.L., and Crawley, J.N. (2011). Automated Three□Chambered Social Approach Task for Mice. Current Protocols in Neuroscience 56. 10.1002/0471142301.ns0826s56.

34. Matas, E., Maisterrena, A., Thabault, M., Balado, E., Francheteau, M., Balbous, A., Galvan, L., and Jaber, M. (2021). Major motor and gait deficits with sexual dimorphism in a Shank3 mutant mouse model. Mol Autism 12, 2. 10.1186/s13229-020-00412-8.

35. Löscher, W. (2002). Basic pharmacology of valproate: a review after 35 years of clinical use for the treatment of epilepsy. CNS Drugs 16, 669–694. 10.2165/00023210-200216100-00003.

36. Emrich, H.M., von Zerssen, D., Kissling, W., Möller, H.J., and Windorfer, A. (1980). Effect of sodium valproate on mania. The GABA-hypothesis of affective disorders. Arch Psychiatr Nervenkr (1970) 229, 1–16. 10.1007/BF00343800.

37. Owens, M.J., and Nemeroff, C.B. (2003). Pharmacology of valproate. Psychopharmacol Bull 37 Suppl 2, 17–24.

38. Phiel, C.J., Zhang, F., Huang, E.Y., Guenther, M.G., Lazar, M.A., and Klein, P.S. (2001). Histone Deacetylase Is a Direct Target of Valproic Acid, a Potent Anticonvulsant, Mood Stabilizer, and Teratogen. Journal of Biological Chemistry 276, 36734–36741. 10.1074/jbc.M101287200.

39. Nau, H., Hauck, R.S., and Ehlers, K. (1991). Valproic acid-induced neural tube defects in mouse and human: aspects of chirality, alternative drug development, pharmacokinetics and possible mechanisms. Pharmacol Toxicol 69, 310–321. 10.1111/j.1600-0773.1991.tb01303.x.

40. Meador, K.J., Baker, G.A., Browning, N., Clayton-Smith, J., Combs-Cantrell, D.T., Cohen, M., Kalayjian, L.A., Kanner, A., Liporace, J.D., Pennell, P.B., et al. (2009). Cognitive function at 3 years of age after fetal exposure to antiepileptic drugs. N Engl J Med 360, 1597–1605. 10.1056/NEJMoa0803531.

41. Locke, J., Shih, W., Kang-Yi, C.D., Caramanico, J., Shingledecker, T., Gibson, J., Frederick, L., and Mandell, D.S. (2019). The impact of implementation support on the use of a social engagement intervention for children with autism in public schools. Autism 23, 834–845. 10.1177/1362361318787802.

42. Gilmore, S., Frederick, L.K., Santillan, L., and Locke, J. (2019). The games they play: Observations of children with autism spectrum disorder on the school playground. Autism 23, 1343–1353. 10.1177/1362361318811987.

