## Supplementary material for "Complex social behaviour during an extended period of time in a valproic acid animal model of autism spectrum disorder": List of abbreviations in figures

**Supplementary information :** List of abbreviations in figures

| **ContactNb** | CNb |
| --- | --- |
| **Oral-oralContactNb** | OOCNb |
| **Oral-genitalContactNb** | OGCNb |
| **SidebysideContactNb** | SSCNb |
| **SidebysideContact,oppositewayNb** | SSCONb |
| **SocialapproachNb** | SANb |
| **GetawayNb** | GANb |
| **ApproachcontactNb** | ACNb |
| **ApproachrearNb** | ARNb |
| **BreakcontactNb** | BCNb |
| **FollowZoneIsolatedNb** | FZINb |
| **Train2Nb** | T2Nb |
| **Group2Nb** | G2Nb |
| **Group3Nb** | G3Nb |
| **Group3breakNb** | G3BNb |
| **Group3makeNb** | G3MNb |
| **Group4breakNb** | G4BNb |
| **Group4makeNb** | G4MNb |
| **MoveisolatedNb** | MINb |
| **MoveincontactNb** | MICNb |
| **Nest3_Nb** | N3_Nb |
| **RearingNb** | RNb |
| **RearisolatedNb** | RINb |
| **RearincontactNb** | RCNb |
| **StopisolatedNb** | SINb |
| **WallJumpNb** | WJNb |
| **WaterZoneNb** | WZNb |
| **totalDistance** | TD |
